# Endothelial Arf6 sustains capillary electrical signaling and cerebral blood flow through PIP_2_ regeneration and activation of Kir2.1 channels

**DOI:** 10.64898/2025.12.23.696233

**Authors:** Maria F. Noterman-Soulinthavong, María Sancho, Saúl Huerta de la Cruz, Michael Yarboro, Maurizio Mandala, Masayo Koide, Nathalie Beaufort, Katalin Todorov-Völgyi, Emma Moreland, David Hill-Eubanks, Martin Dichgans, Mark T. Nelson

## Abstract

Brain capillaries sense neural activity and direct blood flow to active regions—a process termed neurovascular coupling that underlies activity-dependent increases in local perfusion (functional hyperemia). A key contributor to functional hyperemic responses is the capillary endothelial cell (cEC) inward rectifier K^+^ (Kir2.1) channel, which is activated by neuronal activity-derived extracellular K^+^ and initiates vasodilatory electrical signals that propagate through the vascular network. Kir2.1 channel function requires continual production of its lipid cofactor, phosphatidylinositol-4,5-bisphosphate (PIP_2_), and is compromised in mouse models of cerebral small vessel (cSVD). Although decreased PIP_2_ availability is a common feature of cSVD, mechanisms underlying PIP_2_ synthesis remain poorly understood. We hypothesized that Arf6, a small GTPase expressed in cECs that stimulates PIP_2_ production, is critical for this process. Using patch-clamp electrophysiology, we demonstrate that inhibiting Arf6 activity progressively decreased cEC Kir2.1 channel activity. This deficit corresponded to loss of capillary-to-arteriole electrical signaling in isolated vessels and diminished functional hyperemia *in vivo*. Exogenously provided PIP_2_ restored Kir2.1 currents and functional hyperemia after Arf6 inhibition or genetic knockdown. Collectively, our data indicate that cEC Arf6 sustains Kir2.1 activity by maintaining PIP_2_ levels and demonstrate that diminished PIP_2_ synthesis is sufficient to impair functional hyperemia. Furthermore, we identify Arf6 as a mechanistic link between PIP_2_ production and endothelial electrical signaling, highlighting Arf6 as a potential therapeutic target for restoring functional hyperemia.

**Significance Statement:** Active brain regions send electrical signals through capillaries to dilate upstream arterioles and increase blood flow. The resulting activity-dependent increase in local blood flow (functional hyperemia) is mediated through inward-rectifier potassium (Kir2.1) channels. These channels— and hence functional hyperemia—require continuous regeneration of the lipid cofactor PIP_2_ (phosphatidylinositol-4,5-bisphosphate). If PIP_2_ is deficient, electrical signaling fails—a defect characteristic of models of cerebral small vessel disease and Alzheimer’s disease. We identify the small GTPase Arf6 as a key to maintaining PIP_2_ and thus preserving capillary Kir2.1 activity and functional hyperemia. Our findings reveal an important pathway for PIP_2_ homeostasis and position Arf6 as a cornerstone upholding functional hyperemic responses, highlighting Arf6 as a target for restoring cerebral blood flow in disease.

## Introduction

The capillary network comprises the smallest and most abundant vessels in the brain vasculature and serves as the nexus for nutrient and O_2_ exchange. Beyond these passive roles, capillaries detect neuronal activity and initiate hyperemic responses, acting as a ‘sensory web’ to support neurovascular coupling, which links local increases in neuronal activity with increased perfusion (functional hyperemia) ^1,2^. This process depends on electrical signaling mediated by capillary endothelial cells (cECs) through strong inwardly rectifying Kir2.1 potassium (K^+^) channels, which exhibit dual activation by extracellular K^+^ and hyperpolarization. These biophysical properties enable sensing of K^+^ released during neuronal repolarization and regenerative propagation of hyperpolarizing signals between electrically coupled cECs ^2,3^. Thus, cEC Kir2.1 is the molecular lynchpin governing the neuronal activity-triggered retrograde hyperpolarization that travels through successive cECs and ultimately reaches upstream arterioles, where it relaxes smooth muscle cells (SMCs) and dilates arterioles, increasing cerebral blood flow (CBF) to the site of signal initiation ^2,4–6^.

Kir2.1 activity in cECs accounts for a substantial portion of the functional hyperemic response in mice and requires binding of its obligate cofactor, phosphatidylinositol-4,5-bisphosphate (PIP_2_)—a plasma membrane phospholipid that regulates endocytosis, exocytosis, and numerous ion channel functions ^2,7–10^. Similar to the case for other second messengers, PIP_2_ pools are small, constituting <1% of total phospholipids, but highly dynamic, turning over every 20–180 seconds in unstimulated cells ^11,12^. The dynamic aspect of PIP_2_ availability is determined by its metabolism. Phosphorylation of PIP_2_ to PIP_3_ by phosphoinositide 3-kinase (PI3K) or hydrolysis to IP_3_ and diacylglycerol by phospholipase C (PLC) depletes PIP_2_, deactivating ion channels that require it, including cEC Kir2.1 channels ^9,13^. Counteracting these depletion mechanisms are lipid kinases that synthesize PIP_2_, primarily through sequential phosphorylation of phosphatidylinositol (PI) by phosphatidylinositol-4-kinase (PI4K) and phosphatidylinositol-4-phosphate (PIP) by type I phosphatidylinositol-4-phosphate 5-kinase I (PIP5KI) (Figure 1A) ^9^. Unimpeded, the enzymes PI4K and PIP5KI can adaptively increase intracellular PIP_2_ synthesis by up to 40-fold and 500-fold, respectively—a remarkable homeostatic feedback mechanism termed ‘regeneration’ by the Hille group ^9^.

**Figure 1.**
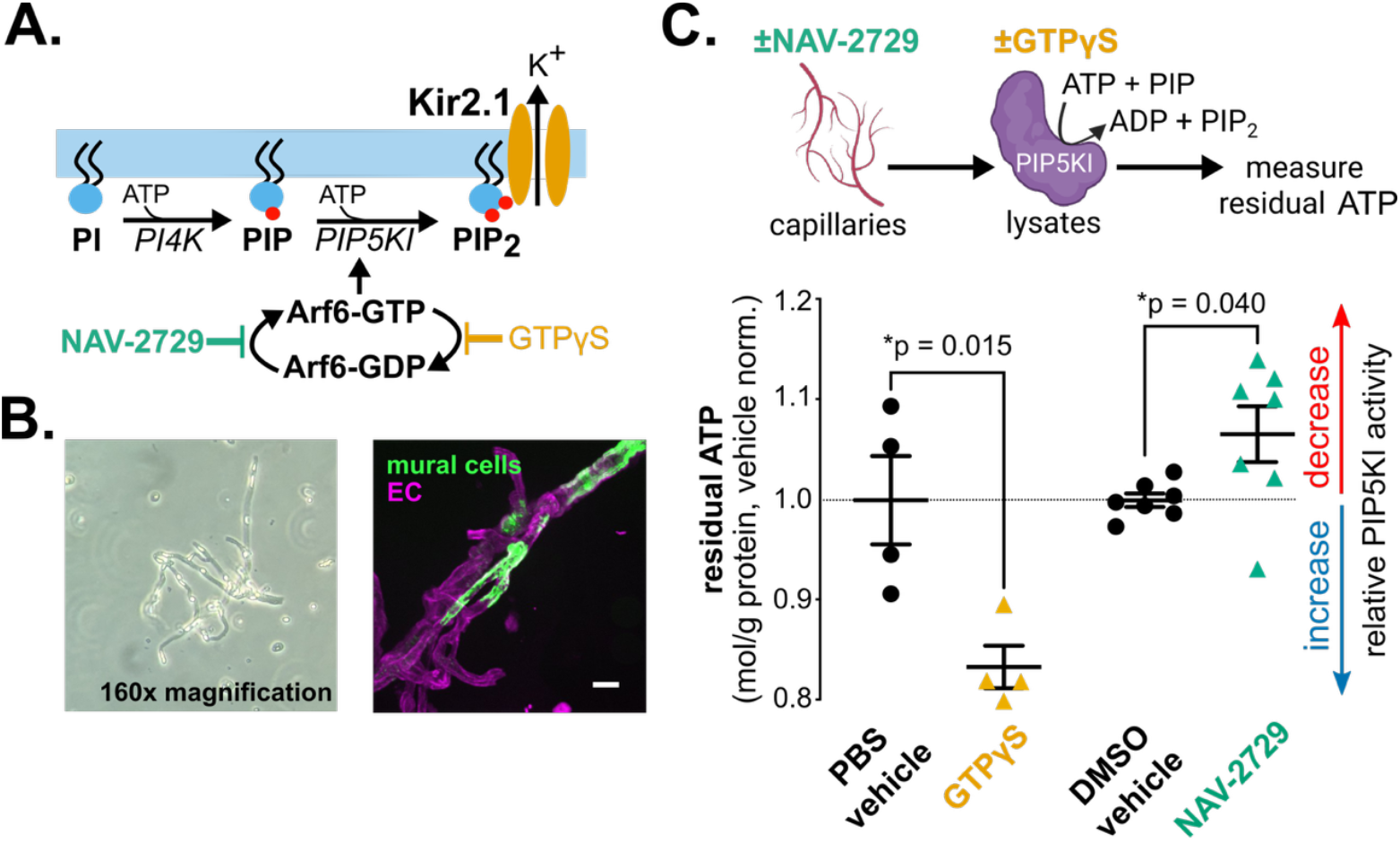
Arf6 regulates PIP5KI activity in brain capillaries. A. Schematic of the predominant pathway for synthesis of PIP_2_ and the proposed regulatory role of Arf6 in brain cECs. NAV-2729 blocks GTP-GDP exchange, promoting the inactive Arf6-GDP form, whereas nonhydrolyzable GTPγS promotes active Arf6-GTP. B. Brightfield microscopy images of isolated brain capillaries from wild-type mice (left), and fluorescence images from mice expressing genetically encoded fluorescent Ca^2+^ indicators in mural cells (right, green) and ECs (right, magenta). Scale bar = 20 µm. C. PIP5KI activity, determined by adding equal amounts of ATP to capillary lysates acutely treated with GTPγS (0.1 mM) or vehicle controls, or NAV-2729 (3 µM) or vehicle controls. Data are normalized to vehicle average per batch (one batch for GTPγS and two for NAV-2729 comparisons) and are presented as means ± SEM (N = 4–7, male C57Bl/6J mice [3–4 months old]; GTPγS comparison by unpaired t-test: t(6) = 3.396; NAV-2729 comparison by unpaired t-test: t(12) = 2.302).

We have shown that inhibiting either PI4K or PIP5KI rapidly impairs cEC Kir2.1 channel activity, suggesting that ongoing PIP_2_ synthesis is essential for sustaining Kir2.1 channel activity ^13^. Furthermore, pathological mechanisms that decrease PIP_2_ synthesis or increase its consumption have been implicated in mouse models of cerebral small vessel disease (cSVD)—the leading cause of vascular dementia ^14–16^. Strikingly, increasing PIP_2_ availability normalizes functional hyperemia in all of these mouse models. Considering the prevalence of vascular dementia and the therapeutic challenge posed by diverse cSVD subtypes, these findings present the appealing concept that multiple cSVD disease mechanisms converge on a common pathway: PIP_2_ metabolism ^17–19^. However, the molecular processes governing PIP_2_ regeneration remain largely unknown ^9^.

To address this gap, we sought to identify candidate cEC proteins that regulate PIP_2_ synthesis and functional hyperemia. Here, we focused on ADP-ribosylation factor 6 (Arf6), a small GTPase that cycles between GTP- and GDP-bound states, a dynamic regulated by the actions of guanine nucleotide exchange factors (GEFs) and GTPase-activating proteins (GAPs) ^20–22^. In its GTP-bound form, Arf6 stimulates PIP_2_ synthesis by enhancing PIP5KI activity at the plasma membrane (Figure 1A) ^23–26^. There is also evidence that Arf6 expression in the cerebrovasculature decreases with age and in CADASIL (Cerebral Autosomal Dominant Arteriopathy with Subcortical Infarcts and Leukoencephalopathy), the most common monogenic cSVD ^27,28^. Yet, while Arf6-mediated PIP_2_ production and PIP_2_-dependent Kir2.1 channel activity are independently well-established, the physiological relationship between Arf6 and Kir2.1 channel activity remains largely uninvestigated. Here, using pharmacological and genetic disruption of Arf6, we demonstrate that Arf6 sustains functional hyperemic responses by maintaining the availability of PIP_2_ for cEC Kir2.1 channels.

## Results

### Endothelial Arf6 stimulates PIP_2_ production sustains Kir2.1 channel activity

Arf6 has many identified roles in ECs ^29–33^, and its primary target, PIP5KI, is expressed in brain cECs as isoforms A–C, all of which share a highly conserved kinase core domain that binds Arf6 ^26,34,35^. To investigate a direct connection between Arf6 and PIP5KI activity in cECs, we measured PIP5KI activity using an ATP depletion kinase activity assay on capillaries from wild-type C57Bl/6J mouse brains ^36^. Included among these isolated capillaries are both ECs and mural cells (Figure 1B). Lysates from capillaries treated with Nav2729 (3 µM) - a small-molecule inhibitor of Arf6-GTP - or vehicle control (0.012% DMSO) were tested for endogenous PIP5KI enzymatic activity by incubating with PIP and ATP. In independent experiments, non-hydrolysable GTPγS (0.1 mM) or its vehicle control (phosphate buffered saline [PBS]) were added to vessel lysates alongside PIP and ATP ^37^. Residual ATP at the end of these incubations was measured as the inverse readout of pan-PIP5KI activity such that greater ATP depletion corresponds to higher PIP5KI activity. GTPγS treatment significantly increased capillary ATP consumption relative to PBS controls, indicating increased PIP5KI activity in the presence of active Arf6-GTP (Figure 1C). Conversely, treatment with NAV-2729 significantly decreased ATP consumption by PIP5KI relative to DMSO controls. Together, these findings indicate that capillary PIP5KI activity in isolated capillaries is sensitive to modulation of Arf6 activity.

We next tested whether Arf6 regulates cEC Kir2.1 channels through PIP_2_, which is required for Kir2.1 channel function ^13^. To this end, we recorded currents from freshly isolated native brain cECs from wild-type mice in the conventional whole-cell configuration in response to 400-ms voltage ramps from −140 to +40 mV (holding potential: −50 mV). Conditions for Kir2.1 activity were optimized by raising extracellular [K^+^]_o_ in the bath to 60 mM with 140 mM K^+^ in the intracellular (pipette) solution ^2^. Under these experimental conditions, K^+^ currents are inward at potentials negative to the K^+^ equilibrium potential (E_K_, −23 mV; calculated by the Nernst equation) and exhibit strong rectification at potentials positive to E_K_. To determine the effects of Arf6 inhibition on Kir2.1 currents over time, we included NAV-2729 (or DMSO) in the pipette to synchronize drug exposure with gaining control over cytosolic content. This design allowed us to analyze currents as a percentage of the initial (t_0_) current amplitude, measured shortly after gaining electrical access to each cell. Importantly, t_0_ Kir2.1 current densities were not significantly different between treatment groups (Supplemental Table 1). To accommodate the relatively slow (over ~15 minutes) loss of Kir2.1 activity following inhibition of PI4K and PIP5K, we performed our voltage ramp protocol every 2 minutes for up to 16 minutes, followed by addition of barium (Ba^2+^)—a selective Kir2 channel pore blocker at the concentration used (100 μM) ^13,38^. For each timepoint, a series of 20 voltage ramps was repeated, and the average maximum inward current at −140 mV was compared between treatments and timepoints.

As experimental treatments, we included 3 μM NAV-2729 (or equivalent volume of DMSO) in the patch pipette solution, with or without inclusion of the water-soluble PIP_2_ analog, diC16-PIP_2_ (10 μM), in the bath. With NAV-2729 in the pipette, Kir2.1 current density steadily decreased over time in cECs, dropping to 49 ± 4% of initial values within our 16-minute recording window (Figure 2A, B). To confirm that the effect of Nav2729 is specific to PIP_2_ availability, we bypassed PIP5KI by bath-applying diC16-PIP_2_, an external source of PIP_2_ that enters cells and maintains Kir2.1 channel activity ^14^. The inclusion of diC16-PIP_2_ in the bath preserved Kir2.1 currents in NAV-2729–treated cECs, maintaining 90 ± 4% of their initial current density. Consistent with our previous studies, Kir2.1 current density in cECs administered vehicle pipette solution remained stable (94 ± 1% of initial) throughout the recordings (Figure 2B, Supplemental Figure 1A) ^13^. These findings suggest that the effects of NAV-2729 on cEC Kir2.1 channel activity are specific to PIP_2_ insufficiency.

**Figure 2.**
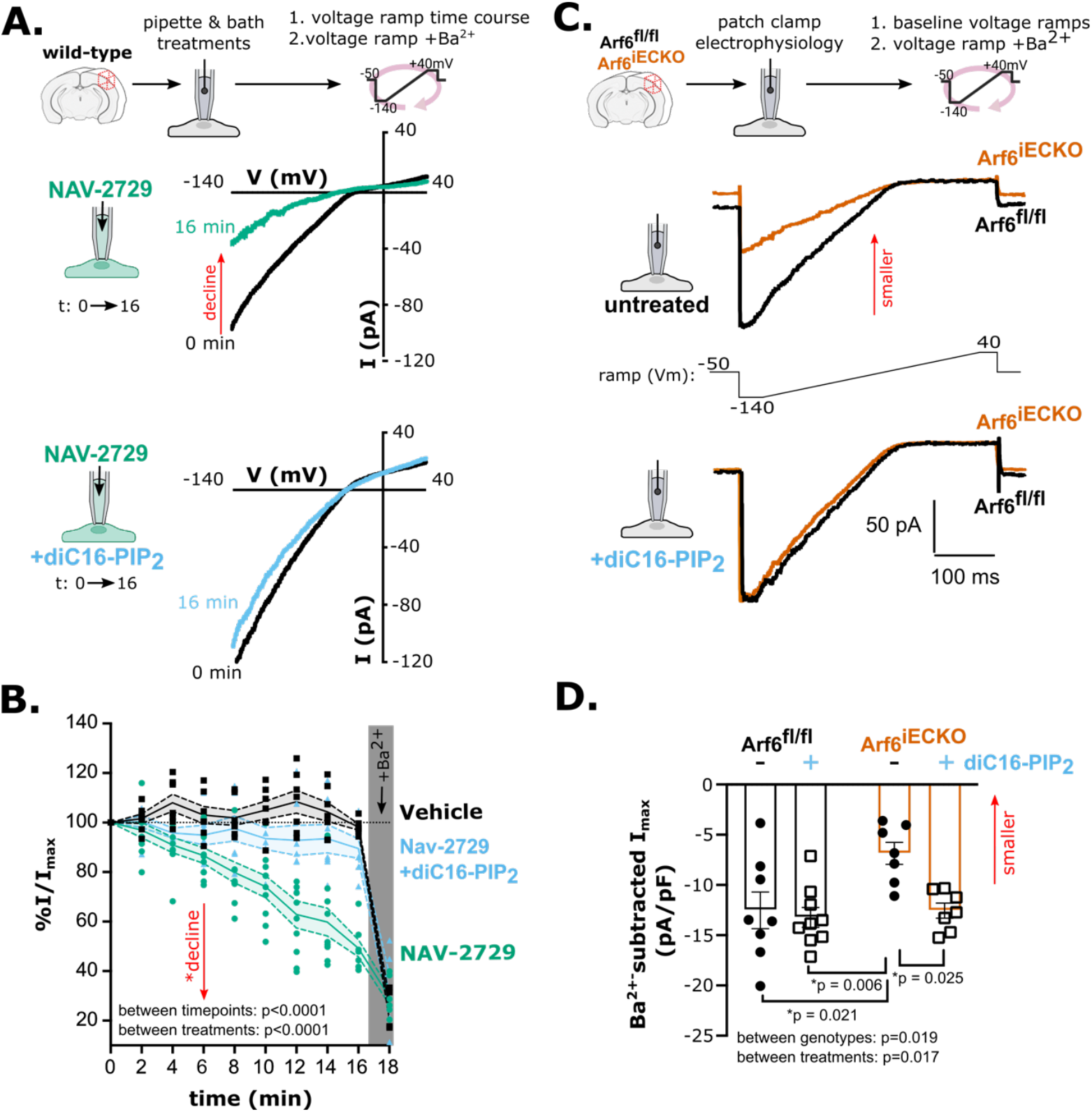
Arf6 controls cEC Kir2.1 channel activity through PIP_2_. A. Experimental strategy (top schematic) and representative traces showing time-dependent changes in Kir2.1 currents (I, pA) in cECs from wild-type C57Bl/6J mice, recorded with NAV-2729 (3 μM; green) in the pipette, without (upper panel) or with (lower panel) diC16-PIP_2_ (10 μM; blue) in the bath. B. Summary data showing the change in maximal inward current for each time point and after final Ba^2+^ treatment, normalized to each cell’s initial current and presented as means (solid line) ± SEM (dashed lines and filled area). Individual cells are shown as single datapoints (n = 6–7 cells per group from 3 male C57Bl/6J mice [3–4 months old]; main effects for time: F(2.39, 46.4) = 85.7, treatment: F(2,21) = 20.64, and interaction: F(4.77, 46.37) = 10.07, *p < 0.0001 for all main effects, mixed effects analysis). Currents in NAV-2729–treated cells were significantly decreased relative to those in vehicle-treated cells for 4–16-minute time points (p < 0.05) and were significantly decreased compared with PIP_2_-supplemented cells for 8–16-minute time points (p < 0.05). Ba^2+^ treatment significantly decreased currents relative to t_0_ for all treatment conditions (Tukey’s multiple comparisons test). C. Representative Ba^2+^-subtracted Kir2.1 currents from Arf6^fl/fl^ (black) and Arf6^iECKO^ (orange) cECs, recorded in the conventional whole-cell configuration, for untreated cells (top) and cells treated with bath-applied diC16-PIP_2_ (bottom). D. Summary data showing effects of bath-applied diC-16-PIP_2_ on Kir2.1 currents in Arf6^fl/fl^ and Arf6^iECKO^ mice. Data are presented as means ± SEM (n = 7–9 cells from 3 mice per treatment condition; two-way ANOVA with Tukey’s multiple comparison post hoc test; interaction main effect: F(1, 27) = 3.8, p = 0.059; genotype main effect F(1, 27) = 6.2); treatment main effect: F(1, 27) = 6.4).

To confirm that the effects of NAV-2729 on PIP_2_ and Kir2.1 currents in cECs are specific to Arf6, we employed our previously described tamoxifen-inducible, EC-specific Arf6-knockout mouse (hereafter, *Arf6*^iECKO^) ^27^. Using the conventional whole-cell patch-clamp configuration on brain cECs from these mice and their wild-type littermate controls, we sequentially performed two series of our voltage-ramp protocol: once under baseline conditions and once following addition of 100 μM Ba^2+^ to the bath. The Ba^2+^-sensitive current was determined (Figure 2C), and the difference in maximum inward current density between genotypes was calculated by normalization to whole-cell capacitance, which is proportional to cell surface area and is unchanged by genotype (10 ± 1 and 10 ± 2 pF in *Arf6*^fl/fl^ [N = 7] and *Arf6*^iECKO^ [N = 8] cells from 3 mice, respectively). Kir2.1 current density was reduced by ~45% in cECs from *Arf6*^iECKO^ mice compared to *Arf6*^fl/fl^ littermates (Figure 2C, D). Bath application of diC16-PIP_2_ restored currents in cECs from *Arf6*^iECKO^ mice without affecting Kir2.1 currents in cECs from *Arf6*^fl/fl^ control littermates (Figure 2C, D). In our 16-minute voltage-ramp time course, NAV-2729 (3 μM) decreased Kir2.1 currents over time in *Arf6*^fl/fl^ littermates (Supplemental Figure 1B, C). In contrast, it had no effect on Kir2.1 currents in cECs from *Arf6*^iECKO^ mice, confirming that Arf6 is the primary molecular target of NAV-2729 at concentrations up to 3 μM, consistent with published reports ^37^. Together, these findings suggest that cEC Arf6 maintains basal PIP_2_ production and sustains Kir2.1 channel activity.

### Arf6 inhibition uncouples capillary-to-arteriole electrical signaling

cEC Kir2.1 channels are essential for sensing neuronal activity and propagating hyperpolarizing electrical signals from capillaries to feeding arterioles ^2^. Our observation that Arf6 inhibition decreased cEC Kir2.1 channel activity suggests that NAV-2729 could impair capillary-to-arteriole signaling and vasodilation. To test whether loss of Kir2.1 channel activity in cECs contributes to vasodilation, we evaluated capillary-to-arteriole electrical signaling and arteriole diameter in 3–4-month-old wild-type mice by *ex vivo* pressure myography using the capillary–parenchymal arteriole (CaPA) preparation, comprising a physiologically pressurized arteriole and intact capillaries from the middle cerebral artery territory (Figure 3A) ^2^. This preparation allows Kir2.1 channel-mediated electrical signal propagation between cECs to be monitored by locally elevating [K^+^]_o_ on capillary extremities (with osmolarity maintained by substituting Na^+^) and measuring changes in the diameter of the upstream, attached arteriole segment. At baseline, all arterioles exhibited basal tone of 33 ± 3%, measured as constriction relative to maximum inner diameter, whether tone was induced by pressure (40 mm Hg) or using the thromboxane A2 analog and G_q_-type G-protein-coupled receptor (G_q_ PCR) agonist, U-46619 (Supplemental Figure 2A; see also Methods) ^39^. Kir2.1-mediated dilation was tested in all preparations by locally picospritzing artificial cerebrospinal fluid (aCSF) containing 10 mM K^+^ onto the capillary bed by pressure ejection (Figure 3A, B). At baseline, all arterioles dilated following treatment of capillaries with 10 mM K^+^; because this was true whether tone was induced by pressure or U-46619, results from both preparations were pooled (Supplemental Figure 2B). Subsequent treatment with NAV-2729 (1 μM) significantly reduced the dilatory response to capillary-applied K^+^ in the attached, upstream arteriole, decreasing arteriole dilation to 28 ± 2% of maximum compared to 49 ± 7% with vehicle treatment (Supplemental Table 2, Figure 3B, C). These data suggest that Arf6 supports cEC-mediated capillary-to-arteriole electrical signaling.

**Figure 3.**
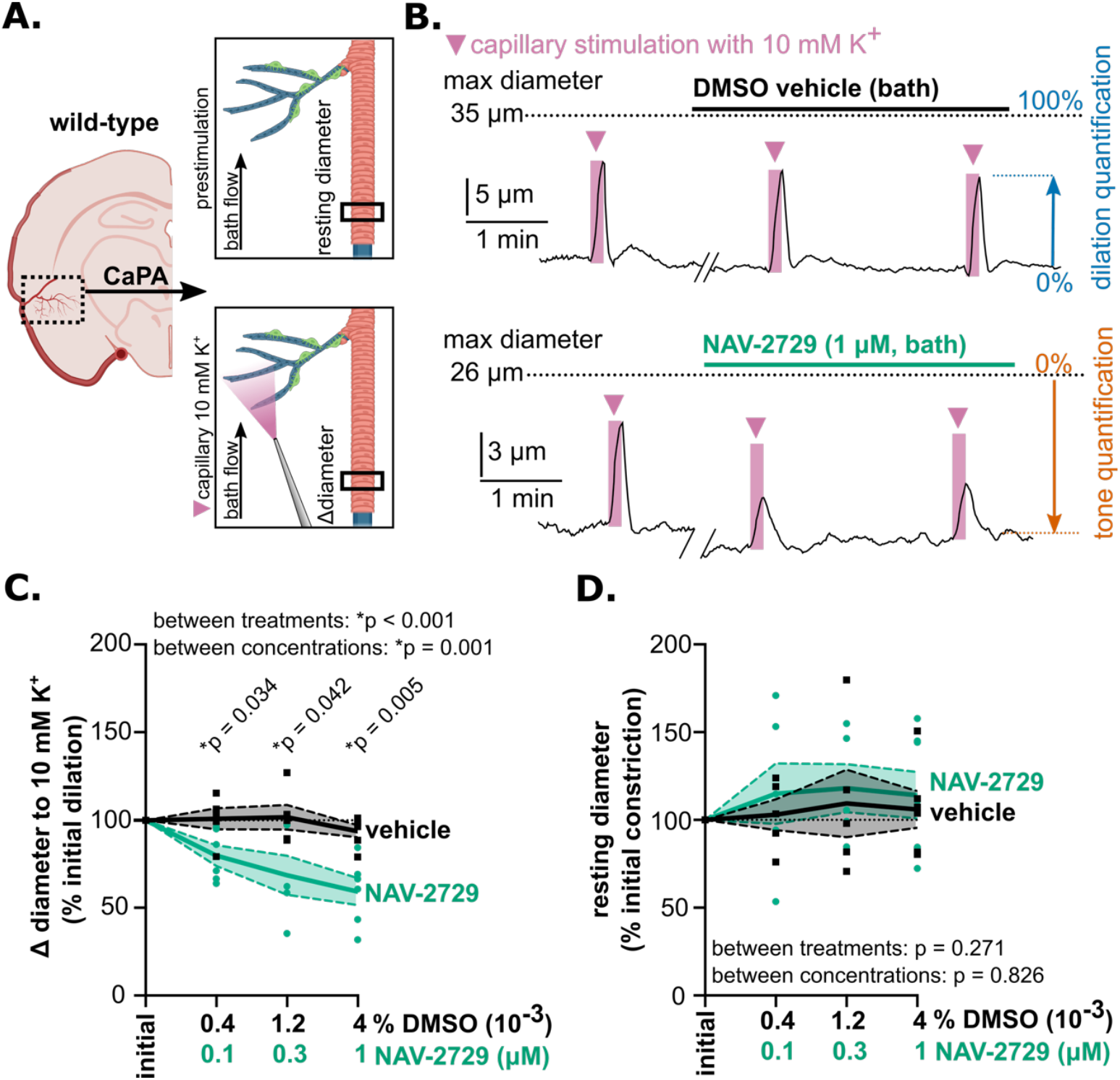
Arf6 inhibition with NAV-2729 decreases arteriole dilation in response to application of 10 mM K^+^ to capillary extremities. A. Schematic of the CaPA preparation for pressure myography measurement of arteriole tone (top) and dilation to capillary-applied 10 mM K^+^ (bottom). B. Representative traces showing arteriole dilation in response to localized picospritzing of capillary extremities with 10 mM K^+^ for 10 seconds under initial conditions and after vehicle (upper trace) or NAV-2729 (1 μM; lower trace) application. C. Summary data showing percent change in arteriole dilation to focal capillary application of 10 mM K^+^, normalized to initial response (*p < 0.05 for vehicle vs. NAV-2729, mixed effects analysis with Tukey’s multiple comparisons test; treatment main effect: F(1,14) = 23.09; concentration main effect: F(2.17, 19.53) = 9.26; interaction main effect: F(2.17, 19.53) = 6.201, p = 0.007). D. Summary data showing percent change in arteriole tone between vehicle and NAV-2729 treatment, normalized to initial tone (p > 0.05; mixed effects analysis; treatment main effect: F(1,14) = 1.31; concentration main effect: F(2.41, 21.67) = 0.241; interaction main effect: F(3, 27) = 0.207, p = 0.891). All data are presented as means ± SEM (N = 5–8 male C57Bl/6J mice [3–4 months old]) and are displayed as individual data points.

Because our quantification of capillary-to-arteriole electrical signaling is dependent on the degree of arteriole constriction, we confirmed that NAV-2729 (0.1–1 μM) did not affect tone in CaPA preparations (Figure 3D, Supplemental Table 3). Additionally, replacing the normal 3 mM K^+^-containing aCSF bath solution with aCSF containing 60 mM K^+^ (osmolarity maintained by substituting Na^+^) caused a similar degree of constriction in arterioles treated with 1 μM NAV-2729 (76 ± 3%) or vehicle (DMSO) control (66 ± 3%), suggesting that inhibiting Arf6 with NAV-2729 does not affect voltage-dependent Ca^2+^ channels or the SMC contractile apparatus (Supplemental Figure 3A, B). During this gradual wash-in of bath-perfused 60 mM K^+^, arterioles transiently dilated before constricting (Supplemental Figure 3A). This dilatory response, which was observed in both NAV-2729–treated (73 ± 4% dilation) and vehicle-treated (59 ± 3% dilation) arterioles (Supplemental Figure 3C), was attributed to the bath solution reaching a K^+^ concentration sufficient to increase Kir2.1 channel activity (7–10 mM) as the 60 mM K^+^ solution mixed with the original 3 mM K^+^ bath solution ^2^. These findings suggest NAV-2729 (0.1–1 μM) does not impair SMC constriction or Kir2.1 function in SMCs or arteriolar ECs.

### EC Arf6 inhibition impairs Kir2.1-mediated functional hyperemia by reducing the availability of PIP_2_

To test whether Arf6-dependent PIP_2_ production sustains Kir2.1-mediated functional hyperemia, we measured changes in CBF in response to whisker stimulation in anesthetized C57Bl/6J mice equipped with an acute cranial window, with and without cortical application of the Arf6 inhibitor, NAV-2729 (Figure 4A). Whisker stimulation-induced CBF responses (i.e., increase from pre-stimulation) following application of NAV-2729 (13 ± 2%) were significantly reduced compared with initial CBF responses in the absence of treatment (23 ± 2%) and CBF responses to vehicle, which were indistinguishable from initial responses in the absence of treatment (Figure 4C, Supplemental Figure 4A). To determine whether these actions of NAV-2729 specifically reflect a reduction in PIP_2_ synthesis, we administered diC16-PIP_2_ intravenously (i.v.), a maneuver that directly delivers PIP_2_ to ECs and restores functional hyperemia in cSVD and Alzheimer’s disease mouse models ^14,16^.

**Figure 4.**
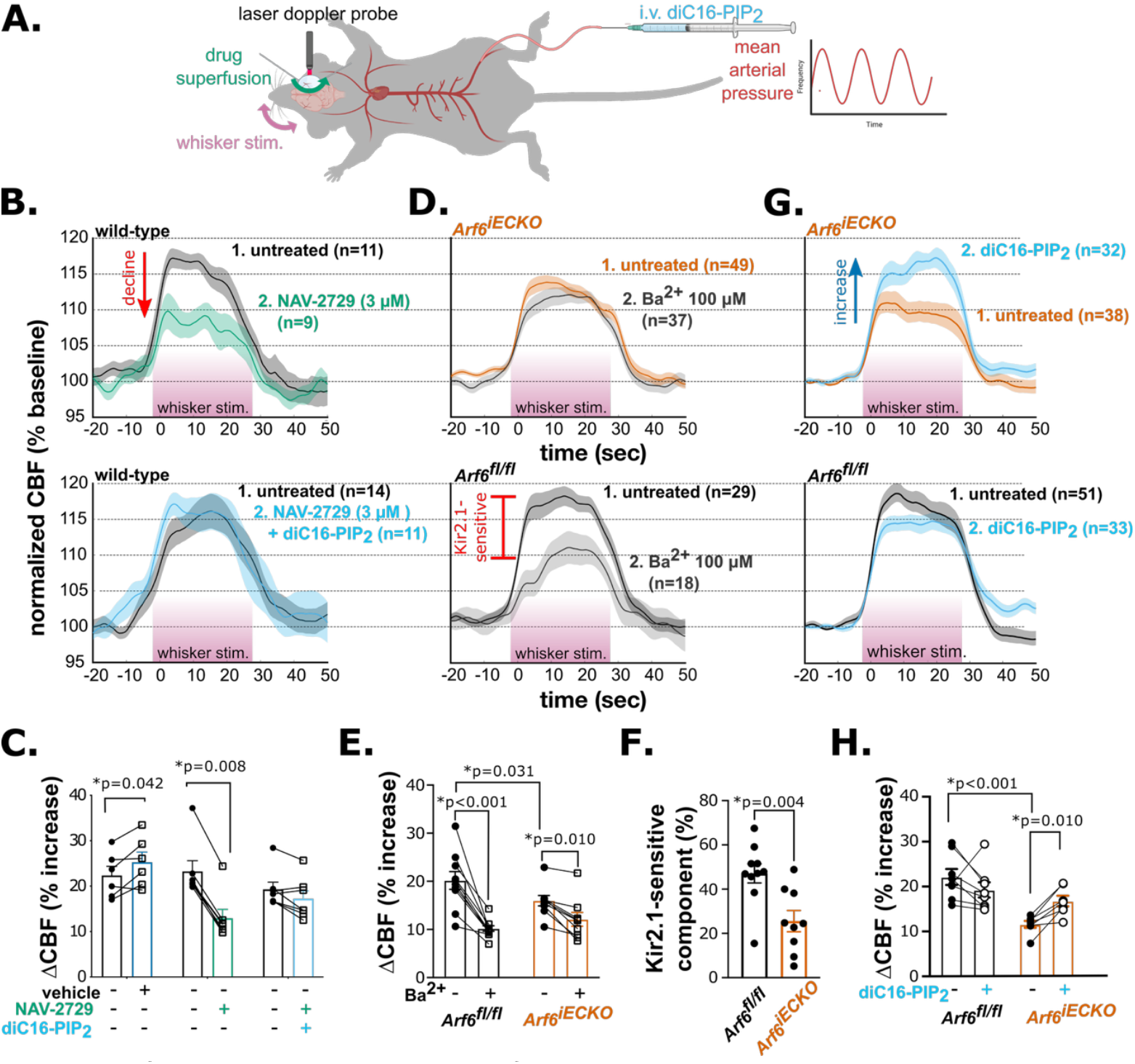
Arf6 loss impairs Kir2.1-mediated functional hyperemic responses to whisker stimulation and PIP_2_. A. Experimental schematic of laser Doppler flowmetry over a somatosensory cortex cranial window. Whisker stimulation (pink) was performed in technical replicates after each treatment. B. Summary CBF traces for sequential recordings obtained at baseline and after cortical superfusion with vehicle, 3 μM NAV-2729, or 3 μM NAV-2729 + 0.5 mg/kg diC16-PIP_2_, normalized to pre-stimulation baseline. Timing of whisker simulation denoted by pink underline. Paired treatments are superimposed. Whisker stimulation technical replicates (n) for all 4–7 animals (N) are shown as means ± SEM. C. Summary data showing paired CBF responses to whisker stimulation under treatment and control conditions. For all panels, each data point represents the average CBF response per treatment per individual animal (N = 4–7 mice; multiple repeated measures t-tests [for comparison between vehicle and untreated baseline; t(5) = 2.713] or Wilcoxon matched-pairs signed rank test; for untreated versus NAV-2729 plus diC16-PIP_2_ comparison, p = 0.781). D. Summary traces showing CBF changes in response to whisker stimulation, before and after Ba^2+^ superfusion (20 minutes), in Arf6^fl/fl^ and Arf6^iECKO^ mice. Whisker stimulation technical replicates (n) for all 9–10 animals (N) are shown as means ± SEM. E. Summary data showing paired CBF responses to whisker stimulation, with and without Ba^2+^ treatment, in Arf6^fl/fl^ (N = 10) and Arf6^iECKO^ (N = 9) mice (repeated-measures, two-way ANOVA with uncorrected Fisher’s LSD post hoc test; genotype main effect: F(1,17) = 0.504, p = 0.487; treatment main effect: F(1, 17) = 56.33, p < 0.0001; interaction main effect: F(1, 17) = 10.91, p = 0.004). F. Summary data showing the Ba^2+^-sensitive component of the total functional hyperemic response (%); same subjects as in D and E (unpaired t-test; t(17) = 3.341). G. Summary traces showing CBF changes in response to whisker stimulation, with and without diC16-PIP_2_ administration (0.5 mg/kg, i.v.), in Arf6^fl/fl^ and Arf6^iECKO^ mice. Whisker stimulation technical replicates (n) for all 7–8 animals (N) are shown as means ± SEM. H. Summary data showing paired CBF responses to whisker stimulation (untreated, plus diC16-PIP_2_) in Arf6^fl/fl^ (N = 8) and Arf6^iECKO^ (N = 7) mice. Data are presented as means ± S.E.M (*p < 0.05, two-way repeated measures ANOVA with uncorrected Fisher’s LSD post hoc test; genotype main effect F(1,13) = 14.47, p = 0.022; treatment main effect: F(1, 13) = 0.9563, p = 0.346; interaction main effect: F(1, 13) = 11.72, p = 0.005).

Applied concurrently with cortical application of NAV-2729, diC16-PIP_2_ supplementation preserved the functional hyperemic response to whisker stimulation (increase from pre-stimulation) compared with initial untreated responses (17 ± 2% vs. 19 ± 2%) (Figure 4B, C), supporting the inference that decreasing Arf6 activity decreases PIP_2_. A lower NAV-2729 concentration (0.3 μM) similarly decreased whisker stimulation-induced CBF responses, and these deficits also were prevented by i.v.-administered diC16-PIP_2_ (Supplemental Figure 4B, C). Collectively, these findings suggest that the NAV-2729-mediated reduction in CBF primarily reflects a decrease in endothelial cell PIP_2_ availability.

To provide further support for the conclusion that the locus of NAV-2729’s impact on functional hyperemia is the vascular endothelium, we measured CBF responses to whisker stimulation in *Arf6*^iECKO^ mice. We found that CBF responses to whisker stimulation (increase from pre-stimulation) were significantly decreased in *Arf6*^iECKO^ mice (16 ± 1%) compared with *Arf6*^fl/fl^ littermates (20 ± 2%) (Figure 4D, E). Moreover, the Kir2.1-sensitive component (relative to total CBF response), determined by subsequent addition of Ba^2+^ (100 μM) to the superfusate, was significantly decreased in *Arf6*^iECKO^ mice (26 ± 5%) compared to *Arf6*^fl/fl^ littermates (47 ± 4%; Figure 4F). Experiments performed in a second cohort of mice reproduced these basal functional hyperemia deficits and further showed that i.v.-administered diC16-PIP_2_ improved CBF responses in *Arf6*^iECKO^ mice compared with initial untreated responses (17 ± 1% vs. 12 ± 1% increase from pre-stimulation). This maneuver yielded CBF responses that were statistically indistinguishable from those in *Arf6*^fl/fl^ littermates (Figure 4G, H), which were unchanged by i.v.-administered diC16-PIP_2_. These data highlight the essential role of EC Arf6 in sustaining PIP_2_ synthesis and Kir2.1 channel function and demonstrate that these actions are crucial for maintaining functional hyperemia.

## Discussion

The brain operates on the brink of energy insufficiency, placing considerable pressure on autoregulatory mechanisms responsible for maintaining basal blood flow and sensitizing neuronal function to metabolic challenges such as diabetic hypoglycemia and insufficient CBF ^40,41^. Layered on top of these constitutive energy requirements are dynamic changes in brain energy demand driven by spatial and temporal variations in neuronal activity ^42^. These latter changes are met by neurovascular coupling mechanisms that elicit functional hyperemia ^43^. Functional hyperemia deficits can be driven by cEC PIP_2_ insufficiency ^14–16^. Yet, the mechanisms of PIP_2_ synthesis remain unexplored in cECs and incompletely understood at large. Using Kir2.1 channel activity as an index for PIP_2_ availability, we show that Arf6 is required to maintain physiological levels of PIP_2_ in cECs. Our data also provide evidence for a direct and causal relationship between EC PIP_2_ homeostasis and CBF regulation, as impairing PIP_2_ metabolism through Arf6 was sufficient to drive functional hyperemia deficits. Collectively, these findings position EC Arf6 as an ‘Achilles heel’ in PIP_2_ synthesis and a new target for treating CBF deficits observed in cSVDs.

The striking effect of Arf6 inhibition on Kir2.1 channel activity suggests that Arf6 controls the rate of PIP5KI activity. Arf6 inhibition decreased cEC Kir2.1 currents by ~50%, and Arf6 knockout decreased Kir2.1 current density by ~45% (Figure 2). By comparison, our previous electrophysiological studies show that precluding PIP_2_ synthesis through PI4K inhibition, PIP5K inhibition, or removal of ATP from the intracellular (pipette) solution causes a ~30–40% decline in cEC Kir2.1 currents ^13^. In other words, inhibiting Arf6 impaired Kir2.1 channel activity to a similar degree as directly inhibiting the enzymes responsible for PIP_2_ production. The strong effect of Arf6 on apparent PIP_2_ availability was initially surprising considering that PIP5KI, the primary target of Arf6 in the PIP_2_ synthesis pathway, is not considered a rate-limiting enzyme ^9^. Indeed, elegant studies by the Hille group showed that the chronically active M-current mediated by the K_V_ 7 channel, which requires PIP_2_, recovers rapidly (t_1/2_ max current within ~10 seconds) following PIP_2_ conversion to PIP by PIP5KI in tsA201 cells ^44^. In contrast, M-current recovery takes ~12 times longer (t_1/2_ max current > 2 minutes) following PLC-mediated PIP_2_ depletion. This prolonged recovery time reflects the required engagement of both PI4K and PIP5KI to replenish PIP_2_ and has defined PI4K as the rate-limiting step for PIP_2_ synthesis. While PI4K is a rate-defining step, our data suggest it is likely not the only one. Indeed, colocalized PI4K and PIP5K synergize PIP_2_ synthesis, underscoring the concept of multiple rate-controlling steps in metabolic pathways ^45^.

In accordance with this notion, endogenous PIP5KI activity varies up to 500-fold depending on its expression level, rate, and localization ^9^. While the mechanisms for this massive dynamic range remain incompletely characterized, our data support a key role for Arf6. Notably, Arf6 can increase PIP5KI activity ~10-fold—and even ~40–60-fold when combined with phosphatidic acid in reconstituted cytosol systems ^23,46^. Additionally, the Arf6 GTP-cycle determines PIP5KI localization at the plasma membrane ^23,24,47^. Finally, Arf6 activation of PIP5KI serves as a negative feedback mechanism against PIP_2_ depletion by PLC (used in the classic Hille kinetic assays) ^9,48,49^. Through these varied mechanisms, Arf6 deletion can disrupt PIP5KI activity and/or localization to block the final step in PIP_2_ synthesis.

Our data also expand on existing models of dynamic PIP_2_ turnover in cECs. Indeed, the classical phenomenon of ion channel activity ‘run down’ due to a mismatch between ongoing PIP_2_ depletion and stunted PIP_2_ synthesis in excised membrane patch recordings has helped to identify many PIP_2_-modulated ion channels ^10,50^. Such continuous PIP_2_ consumption also occurs in native cECs, as illustrated by the observation that impairing PIP_2_ synthesis decreases cEC Kir2.1 channel activity over time ^13^. Notably, none of our electrophysiological recordings involved protocols that drove PIP_2_ depletion; thus, in addition to supporting existing evidence for basal PIP_2_ consumption, our findings introduce Arf6 as a basal regulator of PIP_2_ homeostasis in cECs (Figures 1–3).

While Arf6 is ubiquitously expressed throughout the body ^51,52^, we found that its inhibition specifically impaired Kir2.1 channel activity in capillary beds without affecting K^+^-induced dilation in arterioles (Supplemental Figure 3A, C), suggesting that Arf6 may uniquely control PIP_2_ synthesis and Kir2.1 channel activity in cECs. Interestingly, Arf6 protein abundance progressively decreases with age in cECs but not in mixed capillary cell types, suggesting a cell-type– or vascular-territory– specific vulnerability in Arf6 expression ^27^. On a functional level, loss of Kir2.1 channel activity in cSVD models is also specific to brain cECs; arterial ECs do not show corresponding deficits ^14^. Why cECs might be particularly sensitive to Arf6 regulation of PIP_2_ synthesis remains unknown, but might reflect elevated Arf6 activity at baseline, increased expression and/or activation of Arf6-sensitive PIP5KI isoforms, or some combination of both arising due to increased PIP_2_ turnover and synthesis demands. Emerging evidence suggests that the kinetic and steady state properties of phosphoinositide metabolism vary by cell type, with faster synthesis and larger PIP precursor pool sizes in excitable primary neurons compared with non-excitable primary astrocytes and cultured tsA201 cells ^9,53^. These findings underscore the importance of defining phosphoinositide synthesis kinetics in cECs, which, although not excitable in a classical sense, display abundant electrical and Ca^2+^ signals that directly couple to vasodilatory responses ^10,54^.

Our findings provide the first evidence that Arf6 in brain ECs sustains PIP_2_ levels sufficient for Kir2.1-mediated electrical signaling. However, we did not investigate other ways that Arf6-stimulated PIP_2_ production might control EC function, including through additional PIP_2_-regulated ion channels (e.g., TRPV4, TRPA1, and Piezo1) or through PIP_2_-consuming intracellular signaling cascades (e.g., G_q_ PCRs and PI3K/AKT signaling) ^9,10,55,56^. Hinting at roles for these latter pathways, a recent study by Islam et. al. showed that *Arf6*^iECKO^ mice lose EC-mediated dilatory responses to insulin in vessels from skeletal and adipose tissue but retain G_q_ PCR-mediated dilation to acetylcholine ^32^. These findings suggest that insulin activation couples with Arf6 within the receptor tyrosine kinase and PI3K/AKT signaling cascade. Indeed, others have shown that activation of the receptor tyrosine kinase, Ror2, increases PIP5KI and PIP_2_ at the plasma membrane ^57^. Thus, how Arf6 undergirds different EC signaling pathways through maintenance of PIP_2_ availability warrants further investigation.

From a therapeutic standpoint, most studies on Arf6 have focused on decreasing aberrantly high Arf6 activity in models of cancer, diabetic retinopathy, sepsis, and neurodegeneration ^30,37,58–63^. However, the emerging roles for Arf6 in vasodilatory responses make it clear that any therapy aimed at normalizing Arf6 in disease should do just that—normalize Arf6—no more and no less.

Because PIP_2_ is required for the function of Kir2.1 channels, which underlie electrical signaling-based neurovascular coupling mechanisms, cEC PIP_2_ availability is a major determinant of dynamic fluctuations in CBF. Here, we found that EC Arf6 controls PIP_2_ availability and that pharmacological inhibition or genetic knockdown of Arf6 is sufficient to disrupt EC Kir2.1 channel activity and functional hyperemia. Our findings suggest that endothelial cell PIP_2_ unavailability precipitates functional hyperemia deficits and that Arf6 may thus represent a disease-altering therapeutic target. Accordingly, if PIP_2_ is the ‘coin of the realm’ for ion channel activity, then Arf6 might be the ‘mint’ ^64^.

## Materials and Methods

### Chemicals

diC16-PIP_2_ [PtdIns-(4,5)-P2 (1,2-dipalmitoyl) (sodium salt)] and U-46619 were obtained from Cayman Chemical Company. All other chemicals, including NAV-2729, were obtained from Millipore-Sigma, unless otherwise stated.

### Animal models

All experiments reported in this study were performed at the University of Vermont in accordance with institutional and ARRIVE guidelines and with the approval of the Institutional Animal Care and Use Committee of the University of Vermont. Mice were group-housed on a 12-hour light/dark cycle with *ad libitum* delivery of food and water. Adult (3–4-month-old) male C57/Bl6J mice were acquired from The Jackson Laboratory (Bar Harbor, ME, USA). Dual-reporter mice expressing green fluorescent protein (GFP) in ECs and red fluorescent protein in SMCs and pericytes, hereafter referred to as EC/SMC reporter mice, were created by crossing *Cdh5*-GCaMP8 and *Acta2*-RCaMP mice (both from Jackson Laboratory; CHROMus collaboration) and used to validate capillary purity in independent experiments. EC-specific *Arf6*-KO (*Arf6*^iECKO^) mice were generated by crossing B6.CgArf6tm.11GDP/J mice (*Arf6*^fl/fl^; Jackson 028669) with Tg(*Cdh5*-Cre^ERT2^)1Rha mice (a kind gift from Ralf H. Adams), as described previously ^27,65^. Arf6 deletion was induced by treating *Arf6*^iECKO^ mice with 2 mg tamoxifen/mouse/d over 5 days; *Arf6*^fl/fl^ littermates were treated identically. Mice were used at 3–4 months of age (minimum 1 month of tamoxifen washout). For C57/Bl6J mouse experiments, only male mice were used. These experiments were complemented by experiments in *Arf6*^fl/fl^ and *Arf6*^iECKO^ mice that included both males and females. For experiments with *Arf6*^fl/fl^ and *Arf6*^iECKO^ mice, experimenters were blinded to mouse genotype. There were no differences in bodyweight between *Arf6*^fl/fl^ and *Arf6*^iECKO^ mice (Supplemental Figure 5A). Baseline functional hyperemic responses within each genotype did not differ according to biological sex (Supplemental Figure 5B); thus, males and females were pooled throughout.

### Quantitative polymerase chain reaction (qPCR)

Brain capillary preparations were obtained by density gradient centrifugation using previously reported protocols as described in the Supplemental methods ^36^. *Arf6* gene deletion was determined by qPCR, as described previously ^32^, with modifications as described in the Supplemental methods. Tamoxifen induction (2 mg/mouse/d for 5 days) decreased Arf6 expression in capillaries from *Arf6*^iECKO^ mice by 33.5 ± 6.8% compared with *Arf6*^fl/fl^, results comparable to our previously reported knockout efficiency ^27^.

### PIP5KI activity assay

Brain capillary preparations were obtained by density gradient centrifugation using previously reported protocols as described in the Supplemental methods ^36^. Freshly isolated brain capillaries were resuspended in HEPES-buffered (pH 7.4) DMEM base medium (Agilent, Santa Clara, California, USA) containing 4 mM glucose, 1 mM pyruvate, and 2 mM L-alanyl-L-glutamine. After adding 3 μM NAV-2729 or vehicle (0.01% DMSO), vessel suspensions were incubated for 20 minutes at 37°C with constant mixing. At the end of the incubation, vessels were diluted 1:1 in ice-cold base medium, pelleted by centrifuging at 2,000 x g for 10 minutes at 4°C, and flash frozen. Capillaries were homogenized in ice-cold RIPA buffer (Cell Signaling Technology, Danvers, Massachusetts, USA, 9806S) supplemented with protease and phosphatase inhibitors. Insoluble fractions were removed by centrifugation at 14,000 x g for 10 minutes at 4°C. Equal amounts of capillary protein lysate (estimated by bicinchoninic acid assay) were tested for PIP5KI activity according to instructions provided by the kit manufacturer (K-5700; Echelon Biosciences, Inc., Salt Lake City, Utah, USA). In this assay, ATP is limiting and PIP5KI activity is driven by supplementing capillary lysates with excess PIP substrate and cofactors. At the end of the assay, enzymatic activity is quenched, and PIP5KI activity in the sample is calculated as the inverse of the remaining [ATP], determined by reference to a standard curve, and averaged across technical replicates. To control for batch-to-batch variation, we normalized residual [ATP] to the average vehicle signal per assay. Protein input and incubation time were determined in independent pilot assays (not shown).

### Electrophysiology

Freshly isolated brain cECs were patched in the conventional whole-cell configuration as described previously, with solution compositions and additional details in the Supplemental methods ^2^. For PIP_2_ rescue experiments, 10 μM diC16-PIP_2_ was included in the bath solution, and cells were incubated for 20 minutes prior to patch clamping. The pipette solution contained either 3 μM NAV-2729 or vehicle (0.01% DMSO) ^66^. After gaining electrical access to a single cEC, we performed twenty, 400-ms voltage ramps every 2 minutes for a maximum of 16 minutes. After a 16-minute recording, 100 μM Ba^2+^ was added to the bath to inhibit Kir2 channel activity. Whole-cell currents were normalized to whole-cell capacitance, determined through the cancellation circuitry in the voltage-clamp amplifier.

### Myography

*CaPA preparation:* Parenchymal cerebral arterioles with attached capillary beds were dissected from the middle cerebral artery as previously described and outlined in the Supplemental methods ^2^. Tone was induced by applying physiological pressure (40 mmHg). In a subset of preparations, resting tone was increased by treating with the thromboxane A2 analog, U46619 (200 nM), which constricts vessels to the same degree as pressure (Supplemental Figure 2A). A micropipette filled with aCSF containing 10 mM KCl (substituted for NaCl) was attached to a Pneumatic PicoPump picospritzer (World Precision Instruments, Sarasota, FL,USA) positioned in front of the capillary bed, oriented in the direction of the bath flow. In this configuration, a 10 mM KCl pipette solution pressure-ejected at 10 psi for 10 seconds targets the capillary bed only, so changes in arteriole diameter are attributable to capillary-to-arteriole communication ^2^. Again, both vessels with myogenic tone and U46619-induced preconstriction exhibited the same degree of arteriole dilation to capillary 10 mM K^+^ (Supplemental Figure 2B). Therefore, vessels with myogenic tone and those treated with U46619 were pooled for subsequent analysis. Changes in lumen diameter are presented as the average internal diameter over the last 10 seconds of treatment normalized to the maximal internal diameter in 0 mM Ca^2+^ aCSF (applied at the end of each protocol) using the equation, (average change in diameter)/(maximal diameter) x 100%.

### *In vivo* laser-Doppler flowmetry

Acute cranial windows were generated as previously described and in the Supplemental methods ^2^. Blood pressure was continuously monitored at 200 Hz via the femoral artery cannula and was unchanged between genotypes (Supplemental Figure 5C). Non-fasted blood glucose in blood samples collected at the end of experiments was analyzed using an epoc Reader (Siemens Healthineers, Ottawa Canada) and BGEM test cards (Siemens Healthcare Diagnostics, Dublin, Ireland). No difference was detected between *Arf6*^fl/fl^ and *Arf6*^iECKO^ mice (Supplemental Figure 5D) ^32^. Functional hyperemia was induced by manually brushing the contralateral vibrissae. In controls testing inherent experimenter variability in this manual stimulation paradigm, the same stimulation of the ipsilateral vibrissae induced only a ~5% CBF change (Supplemental Figure 6). If it was not possible to induce an increase in CBF greater than 5% of pre-stimulation levels across the exposed cortical surface, the animal was excluded from further experiments.

Mice were randomly assigned to treatment groups. For acute treatments, NAV-2729 or DMSO vehicle was added to the aCSF superfusate for 20 minutes after performing baseline recordings. Ba^2+^-sensitive whisker stimulation responses were obtained by treating for 20 minutes with Ba^2+^ (100 μM), added to the cortical superfusate. All whisker stimulation responses, recorded in arbitrary units, were analyzed using LabChart software (ADInstruments, Dunedin, New Zealand) and presented as the percent change in CBF from pre-stimulation values, averaged over three whisker stimulation trials per treatment condition. The Ba^2+^-sensitive functional hyperemic response was determined by subtracting functional hyperemia responses under Ba^2+^ treatment from untreated baseline, then multiplying the result by 100%.

### Statistical analysis

Excel was used for data preprocessing, and GraphPad Prism version 10 was used for statistical analyses. Biological outliers were excluded using Grubb’s Test. Data distribution was tested with the Shapiro-Wilk’s normality test. Statistical tests were all two tailed and and are indicated in figure legends. Data are presented as means ± SEM. Differences between means with p-values < 0.05 were considered statistically significant.

## Data Availability

Source data supporting the findings of this study will be available within the paper and its supplementary files.

## Acknowledgments

We thank O. Harraz, L. Schröger, D. Paquet, and J. Gonzalez-Gallego for designing and performing initial pilot experiments, G. Herrera for making the initial MATLAB code for plotting CBF responses, G. Hennig for helpful discussions on data preprocessing, O. Nadeau for editing the manuscript, and S. Tighe, N. Cashen, T. Wellman, and T. Classen for technical assistance. Research reported in this publication was supported by the Leducq Foundation Transatlantic Network of Excellence (International Network of Excellence on Brain Endothelium: A Nexus for Cerebral Small Vessel Disease 22CVD01 BRENDA, to M.D. and M.T.N.), the Totman Medical Research Trust (M.T.N.), and grants from the National Institute of Neurological Disorders and Stroke (R01-NS-110656 and RF1-NS-128963 to M.T.N; R01-NS140868 to M.K, R01-NS-119971 to Nicholas Tsoukias and M.T.N subcontracted), the National Heart Lung and Blood Institute (R35-HL-140027 to M.T.N.), the National Institute of General Medical Sciences (P20-GM-135007 to M.T.N.), The Spanish Ministry of Science and Innovation (PID2023-147925OA-I00 to MS), the American Heart Association (24POST1188081 to SHC), the Deutsche Forschungsgemeinschaft (DFG, German Research Foundation) as part of the Munich Cluster for Systems Neurology (EXC 2145 SyNergy – ID 390857198), CRC 1744 (ID 548585053), DI 722/13-1, and DI 722/21-1; the flagship P4-medicine project DigiMed Bayern; ERA-NET Neuron (MatriSVDs, to MD) and the Vascular Dementia Research Foundation. Technical support for qPCR was provided the Advanced Genome Technologies Core of the University of Vermont and the Larner College of Medicine: Vermont Integrative Genomics Resource Facility (RRID# SCR_021775). Figures Created in BioRender. Noterman, M. (2026): https://BioRender.com/nplbrcw

