## Supplemental Materials for "Endothelial Arf6 sustains capillary electrical signaling and cerebral blood flow through PIP_2_ regeneration and activation of Kir2.1 channels"

**This PDF file includes:**

Supplementary Methods
Supplementary Figures 1 to 6
Supplementary Tables 1 to 3
Supplementary References

### Supporting Information Text

#### Supplemental methods

**Brain capillary isolation.** Whole brains were rapidly dissected and kept in ice-cold media or a chilled centrifuge (4°C) throughout the isolation procedure. Brain capillary preparations were obtained using previously reported protocols [1]. Approximately 90% of vessels isolated in this manner are capillaries (i.e., <10 μm in diameter). Briefly, individual brains were homogenized in a loose-fitting Dounce homogenizer in MCDB 131 media (Thermo Fisher Scientific, Waltham, Massachusetts, USA, #10372019) after first rolling the brain over a Kimwipe to remove meninges and meningeal vessels. The homogenate was centrifuged at 2,000 x g for 5 minutes, after which the resulting pellet was resuspended in a solution of 17.5% dextran in PBS (pH 7.4) and centrifuged at 4,400 x g for 20 minutes with low brake. The resulting myelin plug was removed, and the vessel pellet was gently suspended in PBS and washed over a 40-μm cell strainer. Retained capillaries remaining on the filter were recovered by reverse filtration with 0.5% fatty-acid–free bovine serum albumin (BSA) in MCDB media. Capillaries were pelleted at 2,000 x g for 10 minutes, washed, resuspended in PBS, and centrifuged at 2,000 x g for 10 minutes to yield a final pellet free from media substrates and BSA. Capillaries were kept on ice and used immediately for subsequent assays. Crude yield was determined by capillary wet weight. Isolated capillaries were identifiable based on morphology. Vascular cell content was also visually confirmed using fluorescent EC/SMC reporter mice. Briefly, capillaries were plated on a glass chamber in PBS and imaged using a Revolution spinning-disk confocal microscope equipped with dual iXon Life 16-bit cameras (Andor Technology) attached to a Nikon microscope with a 40X (numerical aperture 0.8) objective. Fluorescence was excited with a 488 nm or 560 nm solid-state laser, and emitted fluorescence was collected through a 527.5/49 nm or 641.5/117 nm band-pass filter. Laser power and exposure settings were established by selecting a vessel expressing both reporters. Remaining vessels were selected at random across the dish and imaged with a rapid serial z-stack. Z-stacks and channels were merged using ImageJ software. Capillaries were used for quantitative polymerase chain reaction and for PIP5KI activity assays.

**Quantitative polymerase chain reaction (qPCR):** *Arf6* gene deletion was determined by qPCR, as described previously [2], with modifications. Briefly, total mRNA was extracted from isolated brain capillaries using a NucleoSpin RNA Plus XS kit (Macherey-Nagel, Düren, Germany) according to the manufacturer's protocols and quantified using a Nanodrop spectrophotometer (ND1000). cDNA was synthesized from 140 ng total mRNA using QuantiTect Reverse Transcription Kit (Qiagen, Hilden, Germany) as described by the manufacturer. qPCR was performed in 96-well plates on an ABI QuantStudio 6 Flex system using SYBR GreenER reagents (Thermo Fisher Scientific) and the standard manufacturer protocol. Gene expression was normalized to peptidylprolyl isomerase A (*Ppia*), and fold-change between $Arf6^{fl/fl}$ and $Arf6^{iECKO}$ capillaries was calculated using the $2^{-DDCt}$ method. Primers were validated by Primer-Blast (NCBI), uniform peak melt curves, amplification product size, and crossing threshold values < 34. Sequences of primer pairs (Integrated DNA Technologies, Inc., Coralville, IA, USA) were as follows: *Arf6*, 5'-ATG GGG AAG GTG CTA TCC AAA ATC-3' (forward) and 5'-GCA GTC CAC TAC GAA GAT GAG ACC-3' (reverse); and *Ppia*, 5'-AGG ATT CAT GTG CCA GGG TG-3' (forward) and 5'-GCC ATC CAG CCA TTC AGT CT-3' (reverse).

**Brain cEC isolation**: A 1-mm-thick brain slice was isolated from one hemisphere of the somatosensory cortex region and homogenized in ice-cold isolation buffer (10 mM HEPES pH 7.3, 5 mM NaCl, 80 mM Na-glutamate, 5.6 mM KCl, 2 mM $MgCl_2$, 4 mM glucose) using a loose-fitting Dounce homogenizer. The homogenate was filtered over a 70-μm nylon mesh, after which retained vascular fragments were reverse filtered with isolation buffer containing 0.5 mg/mL neutral protease (Worthington Biochemical, Lakewood, New Jersey, USA), 0.5 mg/mL elastase (Worthington Biochemical) and 100 μM $CaCl_2$, and incubated at 37°C for 23–25 minutes. Collagenase type I (Worthington Biochemical) was then added to a final concentration of 0.5 mg/mL and incubation was continued for an additional 2 minutes at 37°C. The suspension was filtered over a 70-μm nylon mesh, and the filtrate was resuspended in isolation buffer. Finally, retained capillary fragments were triturated 5–6 times with a fire-polished glass Pasteur pipette. Cells were plated in isolation buffer and allowed to adhere for ~40 minutes before patch clamp electrophysiology experiments.

**Electrophysiology:** Conventional whole-cell electrophysiology was performed on freshly isolated brain cECs as previously described [3]. Filamented pipettes were pulled from borosilicate glass (1.5 mm outer diameter, 1.17 mm inner diameter; Sutter Instruments, Novato, CA, USA) and fire-polished to give a tip resistance of 3–6 MΩ. Currents were amplified using an Axopatch 200A amplifier (Molecular Devices, San Jose, CA, USA), filtered at 1 kHz, digitized at 10 kHz, and analyzed with Clampfit version 9.2 software. Cells were bathed in a solution consisting of 10 mM HEPES (pH 7.4), 80 mM NaCl, 60 mM KCl, 1 mM $MgCl_2$, 2 mM $CaCl_2$, and 4 mM glucose. The pipette solution consisted of 10 mM HEPES, 10 mM NaOH, 11.4 mM KOH, 128.6 mM KCl, 1.1 mM $MgCl_2$, 2.2 mM $CaCl_2$, 5 mM EGTA, 1 mM Mg-ATP (pH 7.2) and either 3 μM NAV-2729 or vehicle (0.01% DMSO) with a net 192 nM free [$Ca^{2+}$] [4].

**Myography**:
*CaPA preparation:* Parenchymal cerebral arterioles were dissected from the middle cerebral artery, taking care to keep the capillary bed intact [3]. Proximal arteriole ends were cannulated on glass micropipettes (1.2 mm outer diameter, 0.69 mm inner diameter; Sutter Instruments) on a pressure myograph (Living Systems Instrumentation, Georgia, VT, USA); distal arteriole ends were occluded with a tie, and capillaries were sealed using downward pressure from a glass micropipette. The arteriole was pressurized to 40 mmHg using a gravity column. Preparations were superfused at 4 mL/min with 36°C artificial cerebrospinal fluid (aCSF; 125 mM NaCl, 3 mM KCl, 26 mM $NaHCO_3$, 1.25 mM $NaH_2PO_4$, 1 mM $MgCl_2$, 4 mM glucose, 2 mM $CaCl_2$) and maintained at pH 7.3 by aerating with 5% $CO_2$, 20% $O_2$, and 75% $N_2$. After experimental treatments, preparations were exposed to bath-applied 60 mM KCl (substituting for NaCl in aCSF) followed by $Ca^{2+}$-free aCSF (substituting $CaCl_2$ for $MgCl_2$ and adding 5 mM EGTA). Preparations that failed to constrict by at least 15% in response to either pressure or U-46619 from their maximum pressurized diameter (determined under $Ca^{2+}$-free aCSF), had no initial dilation to capillary-applied 10 mM KCl (indicating damaged endothelium), or had no endpoint arteriole constriction to bath-applied 60 mM KCl (indicative of damage) were excluded from the analysis.

*Data acquisition:* The luminal diameter of the parenchymal arteriole was acquired in three regions at 15 Hz using a charge-coupled device (CCD) camera and analyzed with IonWizard 6.2 edge-detection software (IonOptix, Westwood, Massachusetts, USA).

***In vivo* laser-Doppler flowmetry**: A surgical plane of anesthesia during procedures was produced with isoflurane (5% induction, 2% maintenance). A cranial window (~2 mm) was made over the somatosensory cortex (AP, -0.9 mm; ML, -3.0 mm relative to bregma) with the dura removed. The exposed window was continuously superfused with aerated (5% $CO_2$, 20% $O_2$, 75% $N_2$ at 37°C) aCSF. Mice were transitioned to an anesthesia regimen of α-chloralose (50 mg/kg) and urethane (750 mg/kg). CBF was recorded through the cranial window with a laser-Doppler probe (PeriMed, Järfälla, Sweden) at 200 Hz and smoothed using the triangular (Bartlett) window method over 1511 samples. Body temperature was maintained at 37°C using a servo-controlled heating pad with a rectal temperature probe. Functional hyperemia was induced by manually brushing the contralateral vibrissae (1-minute or 30-second stimulation at 3–4 Hz) over 2–4 separate trials (technical replicates or 'n' in figures) with at least 1 minute between repeated stimulations. If the CBF recording became unstable at any point in the recording (baseline fluctuation equal to or greater than whisker-stimulation induced fluctuation), all data from that mouse were excluded.
Summary traces of CBF responses to whisker stimulation were generated in MATLAB R2023b. In the process of generating summary data for CBF responses to whisker stimulation, sample noise was reduced by resampling excerpts, including pre-stimulation, stimulation and post-stimulation CBF data, by a factor of 20. Quiescent datapoints were defined as those with values less than the mean plus seven times the standard deviation of values in the lowest 35% of values. The mean plus 2.66-times the standard deviation of quiescent values was used to set the standard deviation for quiescent data (SDQ). Datapoints that exceeded SDQ were labeled as an 'event'. The first event within a 15-second window following whisker stimulation was used to mark the CBF event onset and to subsequently align all data. Trials that failed to meet event threshold used the stimulation

time as a fallback for alignment. Data were then normalized by dividing by mean CBF data over the 90 seconds leading up to the onset of whisker stimulation.

Figures

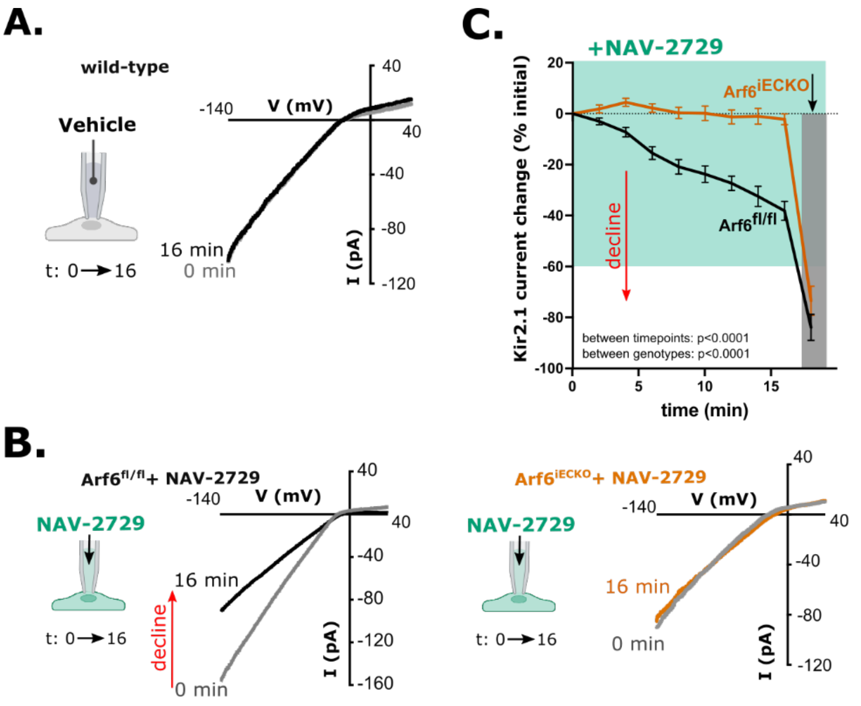


**Supplementary Figure 1.** A. Representative time-dependent changes in Kir2.1 currents (I, pA) in cECs with vehicle (DMSO) in the pipette. B. Representative time-dependent changes in Kir2.1 currents (I, pA) in $Arf6^{fl/fl}$ and $Arf6^{iECKO}$ cECs with NAV-2729 (3 µM; green) in the pipette. C. Time course of the effect of NAV-2729 (3 µM; pipette) on maximal inward current at -140 mV per cell from $Arf6^{fl/fl}$ or $Arf6^{iECKO}$ mice, measured every 2 minutes for 16 minutes. At the end of the time course, residual Kir channel activity was detected (as the difference current) by adding 100 µM $BaCl_2$ (shaded region) to the bath. Data are presented as means ± SEM (n = 5–8 cells per group from five, 3–4-month-old male and female mice; mixed effects analysis with Tukey's multiple comparison test; time main effect: $F(2.434, 30.57) = 182.5$; genotype main effect: $F(1, 14) = 51.17$; interaction main effect $F(9, 113) = 16.13$, $p<0.0001$).

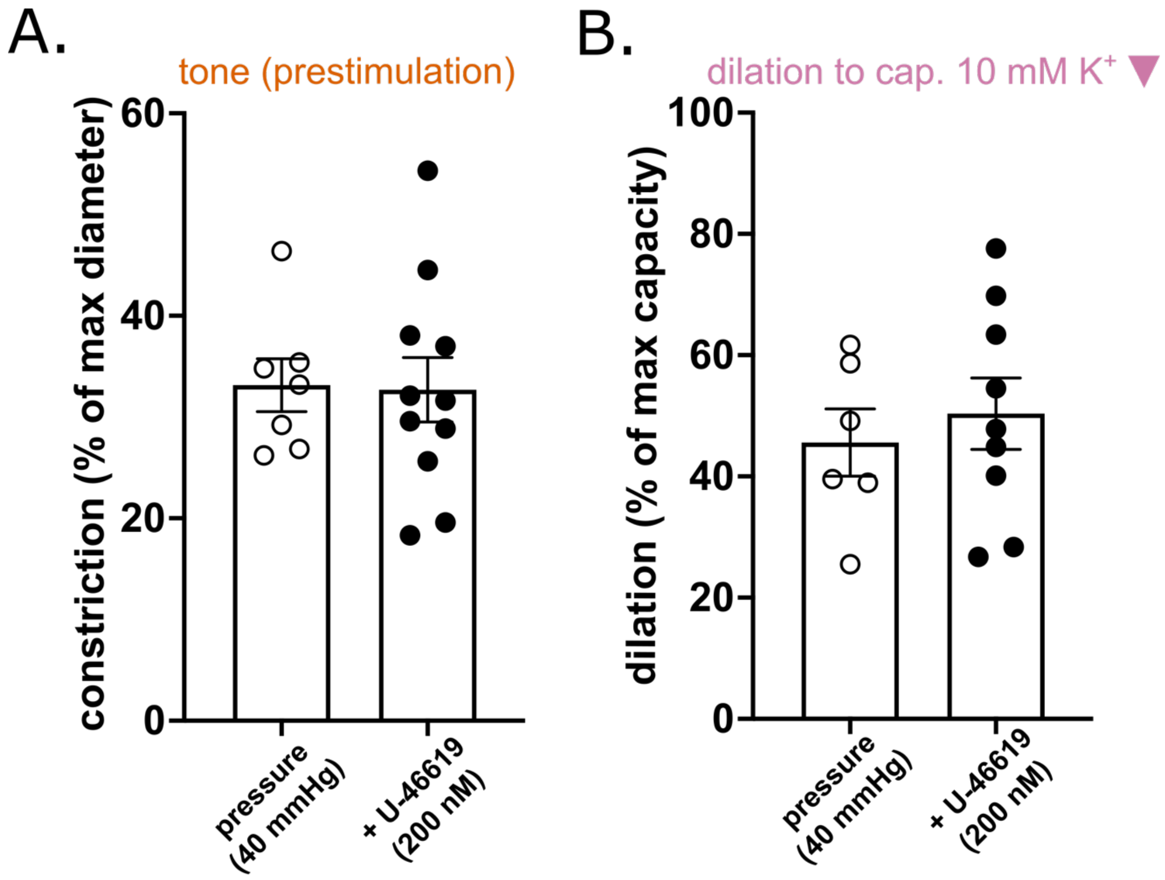


**Supplementary Figure 2.** Tone and dilation to capillary-applied potassium are identical in pressure-induced and pressure- plus U-46619-induced conditions. A. Comparison of tone induced by pressure or pressure (40 mmHg) in addition to the vasoconstricting $G_q$PCR agonist, U-46619 ($t(16) = 0.102$, $p = 0.92$, unpaired t test), and B. corresponding dilations to capillary-applied 10 mM $K^+$ ($t(13) = 0.558$, $p = 0.586$, unpaired t-test). Each datapoint represents an independent preparation from one C57Bl/6J male mouse (3–4 months old). Data are presented as means and SEM (error bars).

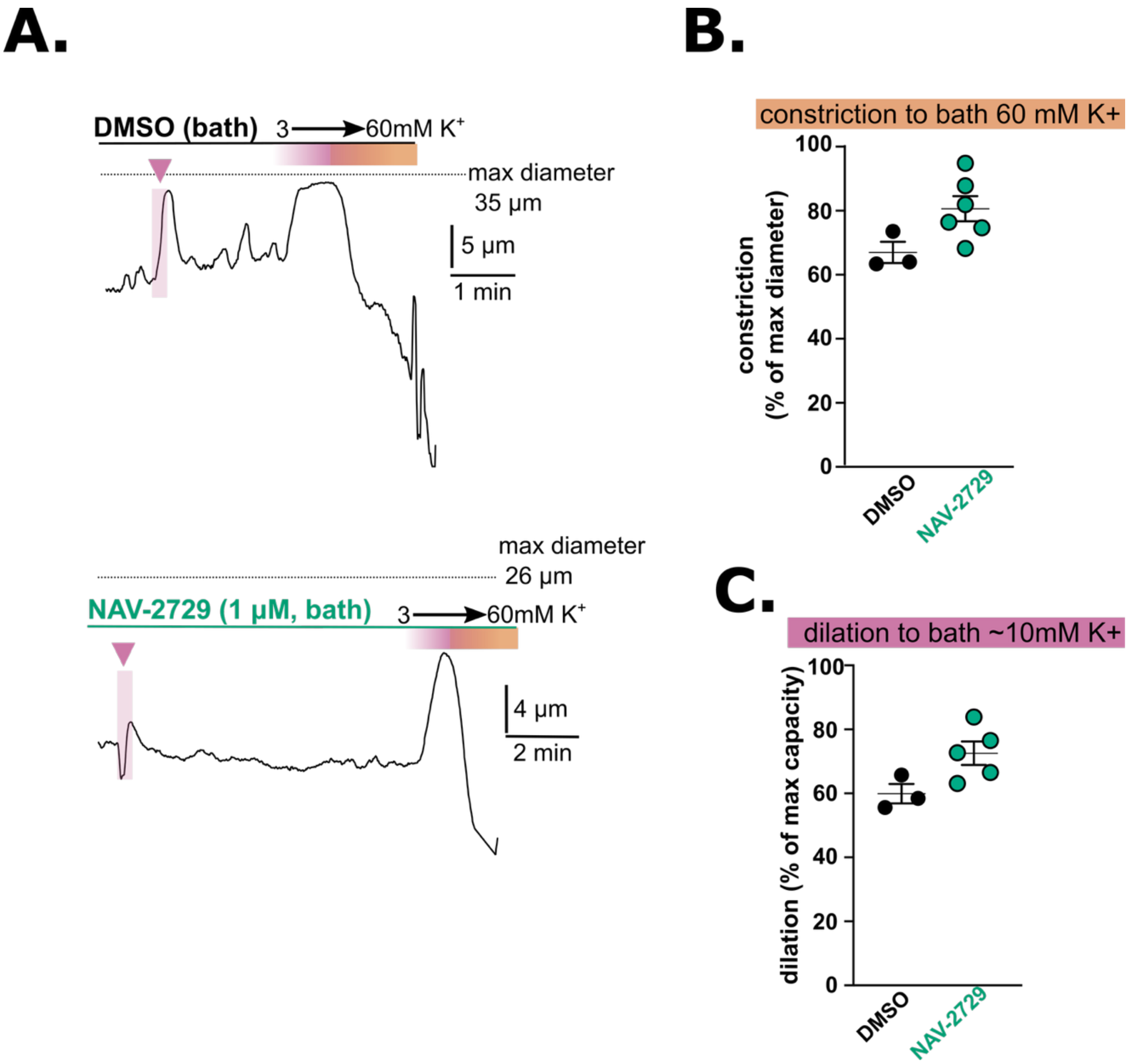


**Supplementary Figure 3.** NAV-2729 (1 μM) does not impair dilation to bath-applied 10 mM $K^+$ or constriction to a depolarizing stimulus. A. Representative traces showing that gradual wash-in of aCSF containing 3 mM $K^+$ to aCSF containing 60 mM K causes transient arteriole dilation followed by constriction in both vehicle- and NAV-2729-treated vessels, while dilation to capillary-applied 10 mM $K^+$ (denoted by pink arrow and shaded region) is decreased by NAV-2729. B, C. Summary data showing maximum constriction to bath-applied 60 mM $K^+$ (B; $t(7) = 2.223$, $p = 0.062$) and maximum dilation to bath-applied elevated $K^+$ (C; $t(6) = 2.355$, $p = 0.057$). Data are presented as means ± SEM (error bars) (n = 3 and 7; unpaired t-test).

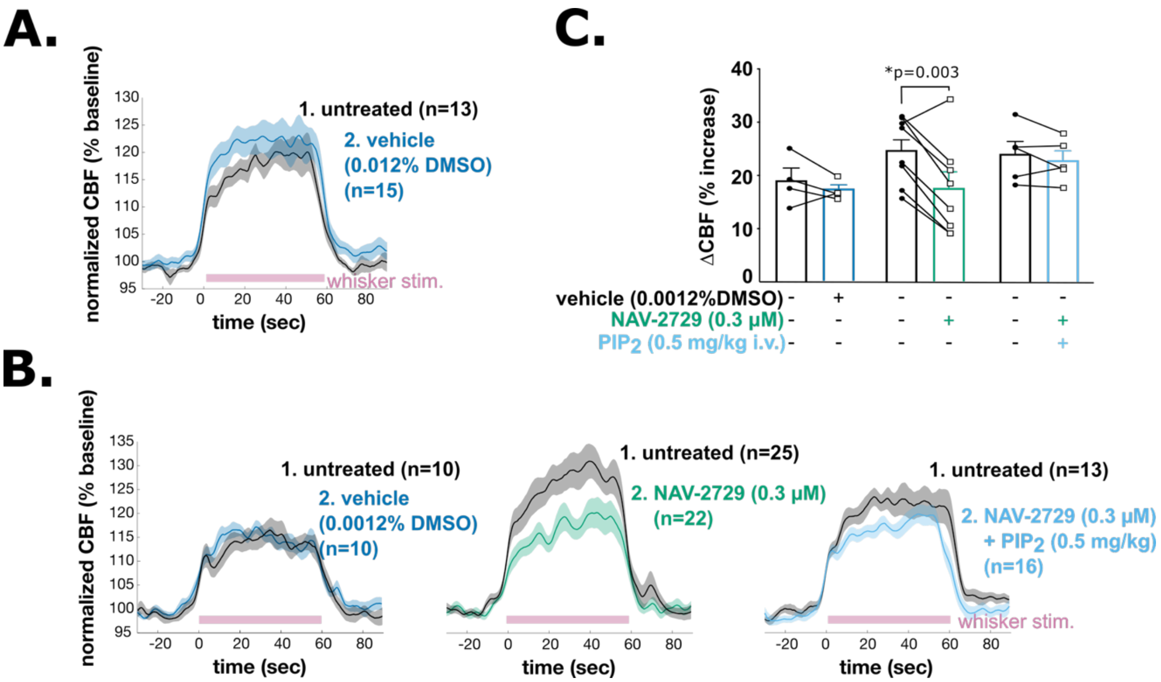


**Supplementary Figure 4**. Low-dose NAV-2729 (0.3 μM) impairs functional hyperemia. A. Normalized CBF traces showing response to 0.012% DMSO treatment (control condition for 3 μM NAV-2729). Technical replicate responses to whisker stimulation (n) were pooled across mice (N = 5) and presented as means ± SEM. B. Normalized CBF traces showing responses to 0.3 μM NAV-2729 treatment and respective controls. Whisker stimulation (pink) was performed in technical replicates after each treatment in the following sequence: 1. Untreated baseline; 2. cortical superfusion with vehicle, 3 μM NAV-2729, or 0.3 μM NAV-2729+0.5 mg/kg diC16-$PIP_2$. Paired treatments are superimposed. Technical replicates across animals (n) are shown as means ± SEM. C. Summary data showing CBF responses to whisker stimulation under baseline and treatment conditions; each data point represents the average CBF response per treatment per individual animal (N = 4–7 mice; *$p < 0.05$, multiple repeated-measures t-tests; for vehicle condition: $t(3) = 1.049$, $p = 0.371$; for NAV-2729 condition: $t(8) = 4.310$; for NAV-2729 plus $PIP_2$ condition: $t(5) = 1.677$, $p = 0.154$).

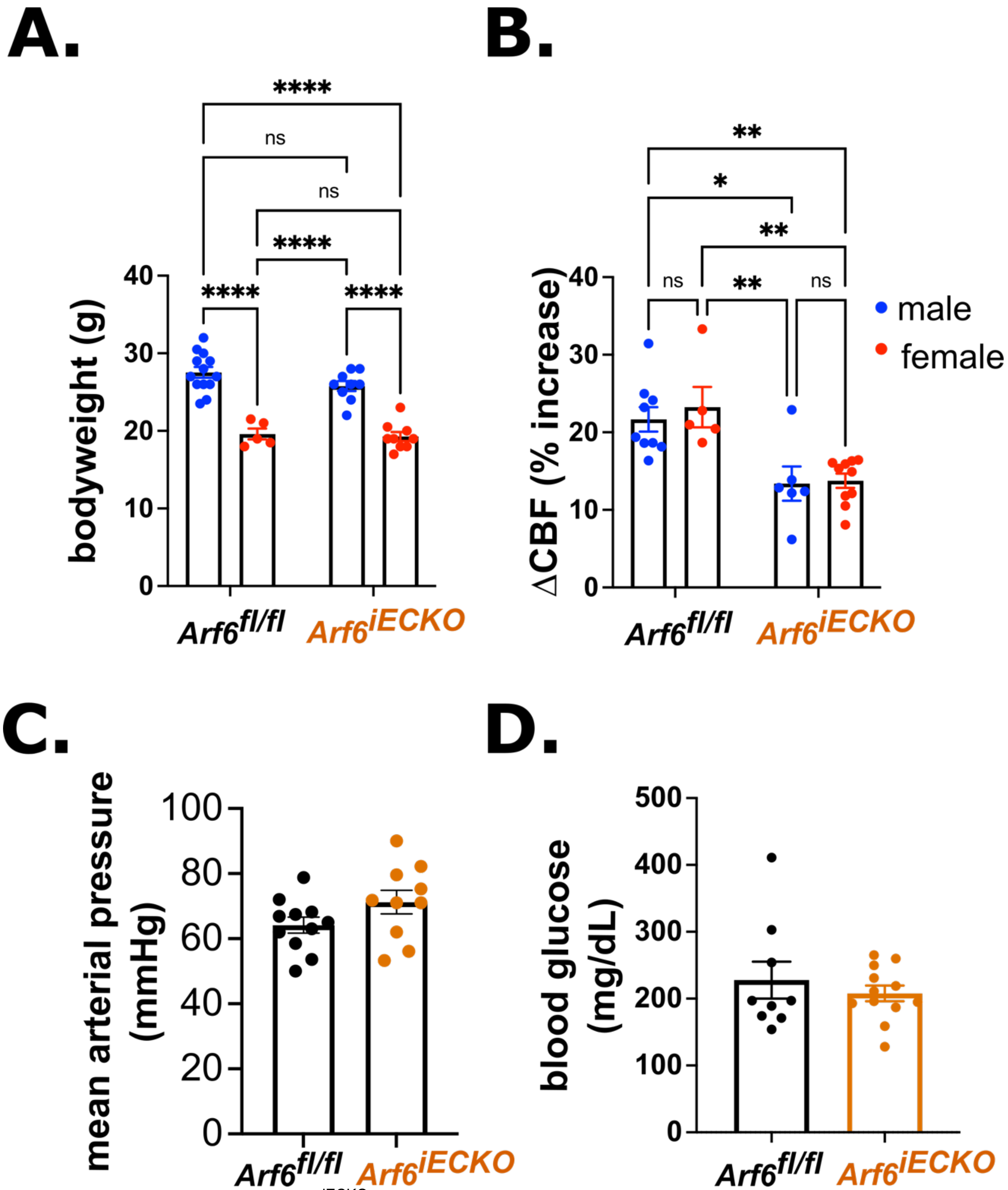


**Supplementary Figure 5.** Arf6$^{iECKO}$ mice do not display sex-specific differences or overt metabolic phenotypes. A. Bodyweight is unchanged in 3–4 month old Arf6iECKO mice compared with Arf6$^{fl/fl}$ littermates (N = 5–13 mice; two-way ANOVA with Śikák's multiple comparisons test; interaction main effect: $F(1,32) = 0.951$, $p = 0.337$; genotype main effect: $F(1,32) = 1.993$, $p = 0.168$; sex main effect: $F(1,32) = 95.78$, $p < 0.0001$). B. Percent CBF increase from baseline in response to whisker stimulation (comparison of baseline data from Figure 4E and H) was unaffected by sex (N = 5–10 mice; two-way ANOVA with Śikák's multiple comparisons test; interaction main effect: $F(1,26) = 0.127$, $p = 0.724$; genotype main effect: $F(1,26) = 26.43$, $p < 0.0001$; sex main effect: $F(1,26) = 0.317$, $p = 0.578$). C. Mean arterial pressure in anesthetized mice at the start of laser Doppler flowmetry recordings is unaffected by genotype (N = 10; p >

0.05, unpaired t-test; $t(19) = 1.642$, $p = 0.117$). D. Non-fasted blood glucose is unchanged in Arf6$^{iECKO}$ mice (N = 9–12 mice; Mann-Whitney test for nonparametric data; $p > 0.999$).

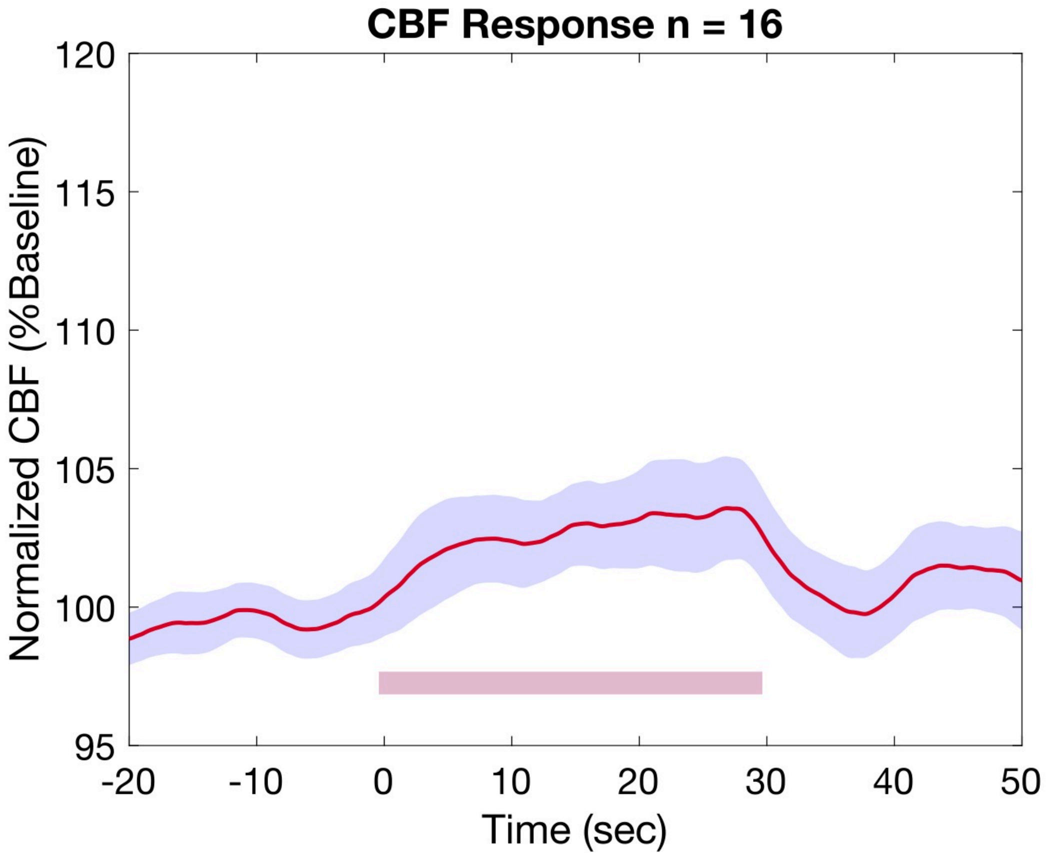


**Supplementary Figure 6.** Background noise in the stimulation paradigm, evaluated by measuring CBF responses to ipsilateral whisker stimulation trials (n) (pink underline) from 12 mice (N).

**Tables**

**Supplementary Table 1.** cEC Kir2.1 channel recording descriptive statistics. Data are presented as means ± SEM, with the number of cells shown in parenthesis. *p < 0.05, **p < 0.01, ***p < 0.001 (mixed-effects analysis followed by Tukey's post hoc test for simple effects between groups [columns])

| | | Vehicle | NAV-2729 | NAV-2729 + diC16-$PIP_2$ |
|---|---|---|---|---|
| Whole-cell capacitance (pF) | | 10.6 ± 0.7 (7) | 9.7 ± 0.4 (10) | 8.8 ± 0.4 (7) |
| Current density (pA/pF) over time (minutes) | 0 | -15.3 ± 2.4 (7) | -14.4 ± 1.6 (10) | -10.4 ± 1.1 (7) |
| | 2 | -15.3 ± 2.1 (7) | -14.1 ± 1.6 (10) | -10.5 ± 1.2 (7) |
| | 4 | -16.4 ± 2.5 (7) | -13.3 ± 1.6 (10) | -9.9 ± 1.0 (7) |
| | 6 | -15.6 ± 2.3 (7) | -12.7 ± 1.6 (10)* | -9.8 ± 1.0 (7) |
| | 8 | -15.5 ± 2.4 (7) | -11.8 ± 1.6 (10)* | -10.1 ± 1.1 (7) |
| | 10 | -15.8 ± 2.1 (7) | -11.0 ± 1.6 (10)** | -9.8 ± 1.2 (7) |
| | 12 | -16.5 ± 2.5 (7) | -9.5 ± 1.6 (10)*** | -9.7 ± 1.3 (7) |
| | 14 | -15.8 ± 2.3 (7) | -9.1 ± 1.6 (9)** | -9.8 ± 1.3 (7) |
| | 16 | -15.4 ± 2.1 (7) | -7.7 ± 1.5 (8)** | -9.3 ± 1.0 (7) |
| | +$Ba^{2+}$ | -3.7 ± 0.6 (4) | -3.3 ± 0.2 (6)** | -3.3 ± 1.1 (4)** |

**Supplementary Table 2.** Arteriole dilation to capillary 10 mM K+, expressed as percent of maximum diameter. Data are presented as means ± SEM, with the number of preparations (equal to number of biological replicates) shown in parenthesis.

| Capillary Treatment | Bath Treatment | Vehicle | NAV-2729 |
|---|---|---|---|
| 10 mM $K^+$ | initial | 50.0 ± 6.8 (7) | 50.2 ± 5.0 (8) |
| | 0.1 μM | 54.1 ± 6.2 (5) | 38.6 ± 3.1 (6) |
| | 0.3 μM | 55.4 ± 7.7 (5) | 32.9 ± 6.6 (5) |
| | 1.0 μM | 48.5 ± 7.4 (6) | 27.5 ± 2.0 (6) |

**Supplementary Table 3.** Arteriole tone expressed as percent of maximum diameter. Data are presented as means ± SEM, with the number of preparations (equal to number of biological replicates) shown in parenthesis.

| Bath Treatment | Vehicle | NAV-2729 |
|---|---|---|
| initial | 27.7 ± 4.0 (7) | 33.2 ± 3.3 (8) |
| 0.1 μM | 30.3 ± 5.4 (5) | 36.8 ± 5.5 (6) |
| 0.3 μM | 30.4 ± 4.7 (5) | 37.9 ± 4.3 (5) |
| 1.0 μM | 28.4 ± 4.4 (6) | 34.0 ± 3.6 (7) |
